# Lipoteichoic acid biosynthesis by *Staphylococcus aureus* is controlled by the MspA protein through competitive interference

**DOI:** 10.1101/2023.05.09.540052

**Authors:** Dora Bonini, Seána Duggan, Alaa Alnahari, Tarcisio Brignoli, Henrik Strahl, Ruth C. Massey

**Affiliations:** School of Cellular and Molecular Medicine, University of Bristol, Bristol, United Kingdom; MRC Centre for Medical Mycology, University of Exeter, Exeter, United Kingdom; Department of Biological Sciences, University of Jeddah, Saudi Arabia; Department of Biosciences, Università degli Studi di Milano, Milan, Italy; Centre for Bacterial Cell Biology, Biosciences Institute, Newcastle University, Newcastle upon Tyne, UK; School of Microbiology, University College Cork, Cork, Ireland; School of Medicine, University College Cork, Cork, Ireland; APC Microbiome Ireland, University College Cork, Cork, Ireland

## Abstract

*Staphylococcus aureus* produces a plethora of virulence factors critical to its ability to establish an infection and cause disease. We have previously characterised a small membrane protein, MspA, which has pleiotropic affects on virulence and contributes to *S. aureus* pathogenicity *in vivo*. Here we report that *mspA* inactivation triggers overaccumulation of the essential cell wall component lipoteichoic acid (LTA) which, in turn, decreases autolytic activity and leads to increased cell size due to a delay in cell separation. We show that MspA directly interacts with the LTA synthesis enzymes UgtP, LtaA and LtaS, and competitively interferes with the association between LtaA and LtaS. While complementation of the *mspA* mutant with wild-type *mspA* reduces the amount of LTA, expression of a mutated version of MspA that does not interfere with the interactions between LtaA and LtaS does not. We suggest that MspA contributes to maintaining a physiological level of LTA in the cell wall by interacting with the LTA synthetic enzymes. In conclusion, this study uncovers the critical role of the MspA protein in regulating cell envelope biosynthesis and pathogenicity.

**IMPORTANCE:** The *S. aureus* cell envelope, comprising of the cytoplasmic membrane, a thick peptidoglycan layer and the anionic polymers lipoteichoic acid and wall teichoic acids, is fundamental for bacterial growth and division, as well as being the main interface between the pathogen and the host. It has become increasingly apparent that the synthesis and turnover of cell envelope components also affect the virulence of *S. aureus*. In this study, we show that MspA, a novel effector of *S. aureus* virulence, contributes to the maintenance of normal levels of lipoteichoic acid in the cell wall, with implications on cell cycle and size. These findings further our understanding of the connections between envelope synthesis and pathogenicity and suggests MspA as a novel a target for therapeutic intervention.

## INTRODUCTION

*Staphylococcus aureus* is a major human opportunistic pathogen (Lowy, 1998), associated with more than 1 million deaths globally in 2019 (Ikuta *et al*., 2022). It colonises asymptomatically 20-30% of the population, which represents a risk factor for later development of disease (von Eiff *et al*., 2001; Wertheim *et al*., 2005). *S. aureus* causes a range of infections from milder skin and soft tissue presentations (SSTIs) such as impetigo to invasive diseases including pneumonia, endocarditis, osteomyelitis and bacteraemia (Tong *et al*., 2015). Key to its ability to cause this range of infections is the production of virulence factors, including adhesins, cytolytic toxins and immune evasins (Foster, 2005; Powers and Wardenburg, 2014). The regulation and utilisation of such factors is complex and is dependent upon the ability of the bacterium to sense and respond to their environment. Critical to all of this is the bacterial cell envelope, which represents the part of the bacteria directly in contact with the host. While maintaining structural integrity and enabling growth and division, the cell envelope is also a main defence barrier to components of the innate immune system such as cationic antimicrobial peptides (AMPs) and fatty acids (Kraus and Peschel, 2008), as well as the target site of clinically relevant antibiotics methicillin, vancomycin and daptomycin (Foster, 2017). As such, the regulation of synthesis and renewal of the envelope components is of great interest both for understanding *S. aureus* pathogenicity and for the development of new therapeutics.

The *S. aureus* envelope comprises the phospholipid bilayer surrounded by the highly cross-linked peptidoglycan sacculus (Turner, Vollmer and Foster, 2014). The cell wall sacculus is interwoven with two types of anionic glycopolymers, wall teichoic acids (WTA), chains of ribitol-phosphate covalently linked to peptidoglycan, and lipoteichoic acids (LTA), chains of glycerol-phosphate anchored to the membrane via a diglucosyl-diacylglycerol (Glc_2_DAG) lipid anchor (Neuhaus and Baddiley, 2003; Barbuti *et al*., 2023). WTA and LTA have overlapping roles in regulating cell division, autolysin activity, cation homeostasis and affinity to AMPs (Weidenmaier and Peschel, 2008). Some of these properties are further regulated by addition of D-alanine residues to the repeating units of both WTA and LTA by proteins encoded by the *dlt* operon, which reduces the overall negative charge of the polymers (Xia, Kohler and Peschel, 2010).

While there are some overlap in their activities, LTA and WTA also perform unique and separate functions (Santa Maria *et al*., 2014). *S. aureus* is viable in laboratory conditions without WTA (D’Elia *et al*., 2006), however, deletion of the LTA synthase *ltaS* is lethal for growth at 37°C, as cells cannot withstand osmotic pressure (Oku *et al*., 2009) and show aberrant septum formation and division (Gründling and Schneewind, 2007b). Deletion mutants of *ltaS* have also been documented to acquire suppressor mutations which bypass the requirement for LTA via different mechanisms such as lowered internal turgor (Corrigan *et al*., 2011; Bæk *et al*., 2016; Karinou *et al*., 2019). The LTA pathway includes GtaB and PgcA that synthesize UDP-glucose, UgtP (also named YpfP) which transfers glucose to diacylglycerol, LtaA which facilitates the flipping of the Glc_2_DAG anchor from the cytoplasmic to the outer leaflet of the membrane (Gründling and Schneewind, 2007a) and LtaS which catalyses the polymerization of glycerol phosphate on the anchor (Gründling and Schneewind, 2007b). Additionally, MprF, which catalyses lysinylation of anionic membrane lipids, has been shown to stimulate LTA synthesis via LtaS in both *Bacillus subtilis* and *S. aureus* (Guyet, Alofi and Daniel, 2023).

We recently identified and characterised a small membrane protein now named MspA (Duggan *et al*., 2020). Inactivation of *mspA* in both the MRSA strain JE2 and the MSSA strain SH1000 was shown to cause a significant decrease in phenol soluble modulins (PSMs) toxin production and cytotoxicity due to downregulation of the accessory gene regulatory (Agr) system. The *mspA* mutant was shown to have a reduced content of the membrane carotenoid pigment staphyloxanthin, to be more susceptible to components of the innate immune response such as fatty acids and AMPs, and to have decreased membrane stability when challenged with the detergent SDS. Interestingly, both systems for uptake (IsdC, IsdB and FhuC) and export (HrtAB) of haem, the major iron source for *S. aureus* during infection, were more abundant in the *mspA* mutant compared to the wild type. The mutant also had increased levels of intracellular iron, suggesting dysregulation of iron homeostasis. These pleiotropic effects observed *in vitro* resulted in attenuation of the mutant in superficial and systemic infection mouse models (Duggan *et al*., 2020).

Given the predicted location of MspA and the pleiotropic effects its inactivation has on virulence and pathogenicity, we hypothesised that MspA could have a structural role, contributing to the synthesis of the membrane or cell wall, or supporting the stability of proteins involved in such processes. Therefore, virulence defects showed by the mutant could be a downstream consequence of perturbations in cellular structural integrity. To test this, here this we show that inactivation of *mspA* an increase in cell size and a delay in cell cycle progression due to overaccumulation of LTA. We demonstrate that MspA interacts with the LTA enzymes UgtP, LtaA and LtaS and suggest that MspA maintains physiological LTA levels in the cell wall by competitively interfering with the ability of these enzymes to interact and synthesise LTA. With such a critical role in cell envelope biosynthesis, and subsequently in pathogenesis, we propose that the MspA protein represents a major target for future therapeutic intervention.

## RESULTS

### The MspA protein affects cell size and cell cycle progression

The *S. aureus* small membrane protein MspA has been shown to have a large impact on virulence, pathogenicity as well as membrane stability (Duggan *et al*., 2020). As it is largely embedded in the membrane and lacks any annotated functional domains, we hypothesised that it could have a structural role and support envelope synthesis directly, or stabilise proteins involved in membrane and cell wall biogenesis. To test this, we first investigated if inactivation of *mspA* caused visible alterations to the membrane or cell wall by performing transmission electron microscopy (TEM). Our first observation was that many of the MspA-deficient cells were noticeably larger than wild type cells (Fig. 1A), which we confirmed by quantifying the cell area as a proxy for cell size (Fig. 1B). While there were no major differences in the envelope architecture between most of the wild-type and mutant cells (Fig. 1A), a small number of *mspA* mutant cells (5 out of >300) had irregular septa which formed before daughter cell separation and were not oriented perpendicular to the previous division plane (Fig. S1). We did not observe these abnormalities in the wild-type population, having examined an equivalent number of cells. To assess whether the increase in cell size was observable throughout the cell cycle or if it was specific to a particular stage, we classified cells in three groups: (i) non-dividing cells without visible septum, (ii) actively dividing cells with incomplete septa, of (iii) divided cells with fully formed septum. The cell size of the *mspA* mutant was significantly increased in all cell cycle stages compared to the wild type (Fig. 1B). We also noticed that the proportion of cells in each of the cell cycle stages varied between wild type and the mutant (Fig. 1D). In particular, the proportion cells with a fully formed septum was more than double for the MspA-deficient strain (48.9% compared to 21.5% of the wild type cells analysed). The fraction of cells in each division stage is proportional to the time that cells spend in that stage (Monteiro *et al*., 2015). Therefore, a higher proportion of cells with fully formed septa suggests that lack of MspA triggers a delay in daughter cell separation after septum formation. It has been shown that *S. aureus* cells with a complete septum elongate before splitting into two daughter cells (Monteiro *et al*., 2015). Therefore, the lengthening of this division stage likely drives the increase in cell size observed in the mutant, while in a small number of cases (Suppl. Fig 1) synthesis of new septum is initiated before daughter cells separation is completed.

**Figure 1.**
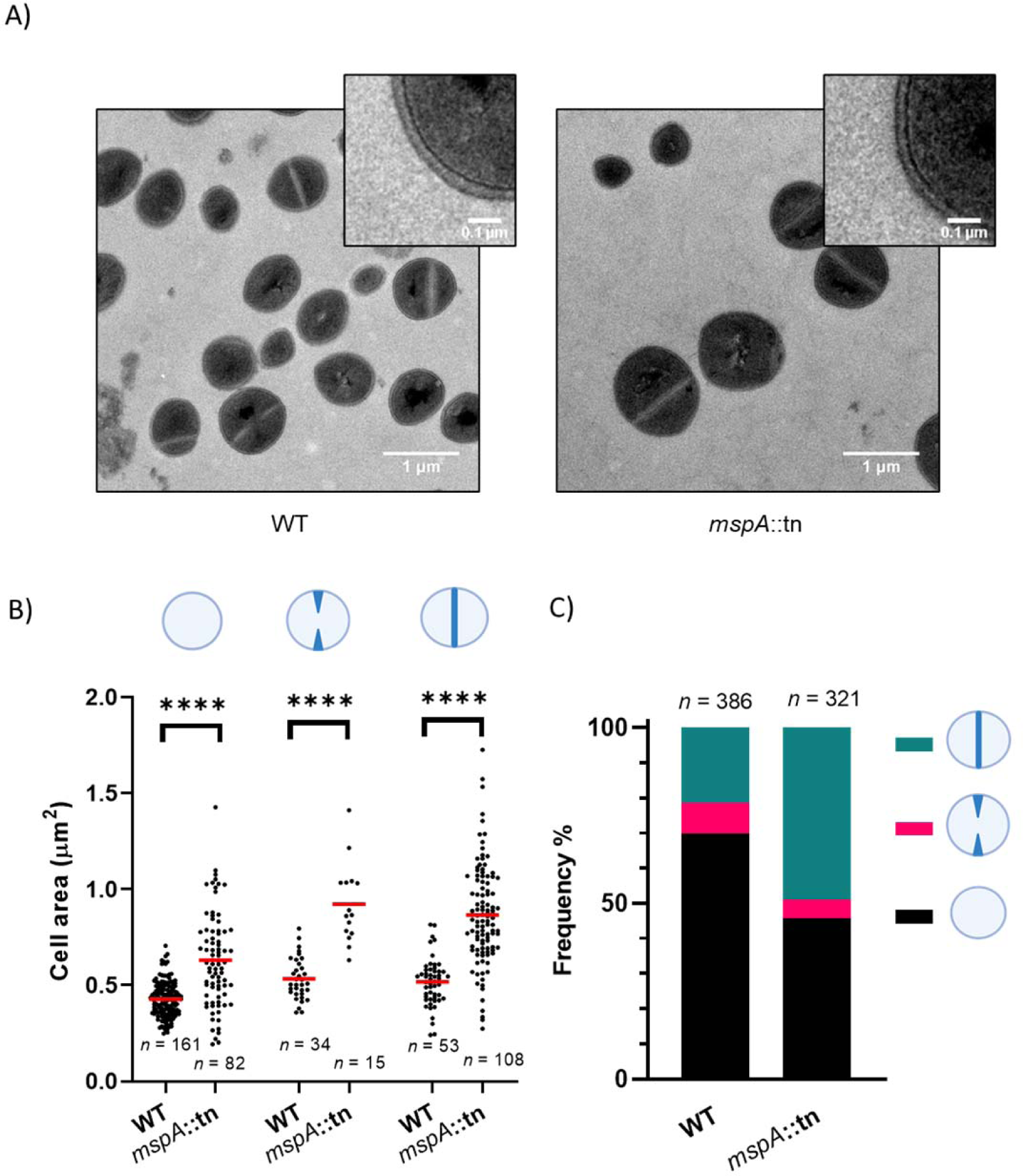
MspA-deficient cells are larger and cell separation is delayed. **A)** Transmission electron microscopy micrographs of SH1000 wild type (WT) and isogenic *mspA* transposon mutant cells. **B)** Measurements of cell area of wild-type and *mspA*::tn cells in each cell division stage. From left to right: non-dividing cells (Welch’s *t*-test, ****, P < 0.0001); dividing cells with an incomplete septum (Welch’s *t*-test, ****, P < 0.0001); dividing cells with a closed septum (Welch’s *t*-test, ****, P < 0.0001). Red lines indicate the mean. C) Frequency of cells in the three phases of cell division. The total number of analysed cells is indicated on top of the stack graph.

### The increase in cell size of the *mspA* mutant is associated with increased abundance of Lipoteichoic acid

A lack of proteins involved in cell division or cell wall synthesis frequently causes an increase in cell size (Cheng, Missiakas and Schneewind, 2014; Do *et al*., 2020; Tinajero-Trejo *et al*., 2022; Wacnik *et al*., 2022). For example, deletion mutants of the genes encoding for UgtP or LtaA, proteins responsible for the lipoteichoic acid (LTA) glycolipid anchor synthesis and flipping (Gründling and Schneewind, 2007a) display longer LTA structures, enlargement of cell size, defects in septa formation and higher susceptibility to lytic enzymes (Hesser, Schaefer, *et al*., 2020). Given the overlap between these phenotypes and those of the MspA-deficient cells described above, and the fact that LTA is anchored to the membrane and spans the membrane proximal region of the cell wall (Reichmann *et al*., 2014; Zhang *et al*., 2021), we hypothesised that the loss of MspA may affect LTA biosynthesis. To test this, we first examined whether lipoteichoic acid abundance was altered in the MspA-deficient cells. Western blots using anti-LTA antibodies (Fig. 2A, Fig. S2) and their quantification (Fig. 2B) showed that MspA-deficient cells grown overnight had significantly increased LTA-content compared to the wild type. Based on their migration, LTA polymers looked of similar molecular weight in both strains and did not display any clear differences in length.

**Figure 2.**
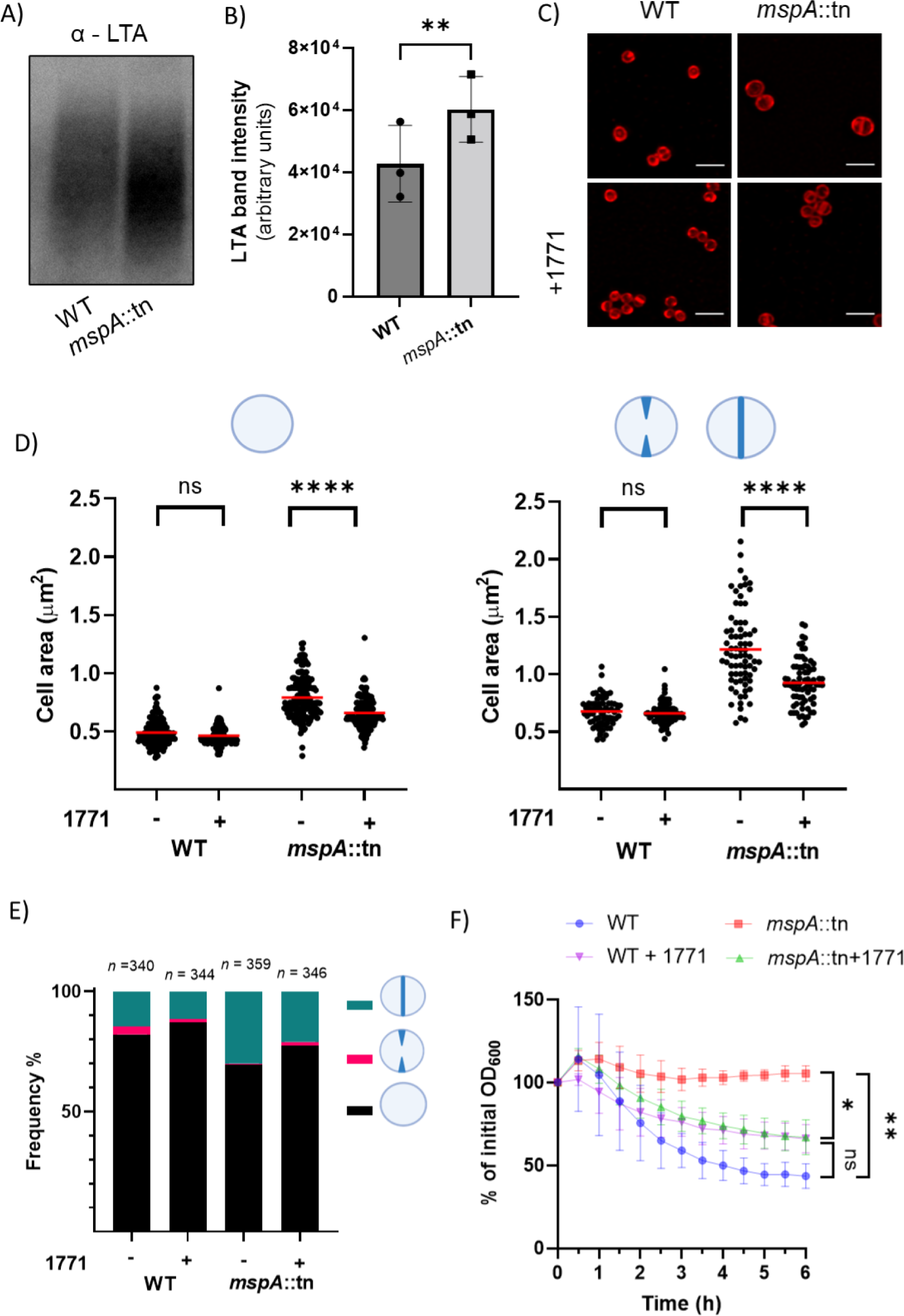
MspA-deficient cells feature an increased lipoteichoic acid content associated with altered cell size and cell cycle progression. A) Western blot with anti-LTA antibody on samples from SH1000 wild type (WT) and SH1000 *mspA*::tn. B) Quantification of the intensity of LTA western blot bands performed with ImageJ (*n*= 3, paired *t*-test, **, P = 0.0045) C) Confocal fluorescent microscopy images of fixed wild-type and *mspA*::tn cells grown in the presence and absence of compound 1771 (2 µg/ml) and stained with Nile Red membrane dye. Scale bar: 2 µm. D) Quantification of the cell area of Nile Red stained cells grown in the presence and absence of compound 1771 (2 µg/ml). 1771 does not affect the wild type cell area of non-dividing cells (left) and dividing cells (right) while both areas are significantly decreased in the absence of MspA. Data were analysed with a two-way ANOVA with Sidak’s *post-hoc* test. Red lines indicate the mean. Left: *n* = 168, untreated vs 1771, WT, ns, P = 0.1; *mspA*::tn, ****, P<0.0001. Right: *n* = 168, untreated vs 1771, WT, ns, P = 0.8; *mspA*::tn, ****, P<0.0001. E) Frequency of cells stained with Nile Red in each of the division stages when grown in the presence and absence of 1771 (2 µg/ml). The 1771 treatment reduces by ∼21% (wild type) and by ∼30% (MspA-deficient) the proportion of cells with a mature septum. Total number of cells analysed is indicated on top of the bars. F) Triton X-100 triggered lysis of wild-type and *mspA*::tn strains grown to exponential phase in the presence and absence of compound 1771 (2 µg/ml), as measured by monitoring OD_600_ over six hours post exposure to Triton X-100 (n = 3, two - way ANOVA with Tukey’s *post-hoc* test; at 6hrs; WT vs *mspA*::tn, **, P = 0.02; WT vs WT + 1771, ns, P = 0.09; *mspA*::tn vs *mspA*::tn 1771, *, P = 0.04).

To test that the increase in cell size of the MspA-deficient cells was due to the altered amount of LTA, we cultured the wild-type and *mspA* mutant overnight with the compound 1771.This has been previously shown to decrease LTA synthesis (Richter *et al*., 2013), although the target of inhibition is as yet unclear (Vickery *et al*., 2018). Since LTA is essential, we chose a sub-growth inhibitory concentration of the compound (2 µg/ml) (Fig. S3, S4) that did not alter OD_600_ after overnight growth. Using Nile Red membrane stain and confocal microscopy, we found that 1771 treatment did not affect cell integrity and there was no lysis observable (Fig. 2B). Measurements of cell area of Nile Red stained cells grown in the absence of 1771 confirmed the enlarged cell phenotype of the MspA-deficient cells observed with TEM. In both non-dividing (Fig. 2D) and dividing cells (Fig. 2E), the 1771 treatment significantly decreased the size of MspA-deficient cells, while it did not affect wild-type cell size. This suggests that the higher content of LTA observed for the MspA-deficient cells is linked to the observed increase in cell size. In wild-type cells producing normal levels of LTA, reducing the polymer synthesis with 1771 does not decrease cell size, possibly as the lower limit in cell size is not as impacted by the abundance of these teichoic acids.

### The greater abundance of LTA in MspA-deficient cells affects the cell cycle by supressing autolysis

Given the association of increased LTA abundance and cell size, we next tested whether the elevated LTA-levels were also responsible for the higher proportion of mutant cells with a mature septum. We cultured cells with and without the 1771 LTA-inhibiting compound as above, stained them with Nile Red, and quantified the prevalence of cells in the different cell cycle stages. This analysis confirmed that MspA-deficient cells exhibited a higher proportion of cells with a mature, closed septum compared to the wild type (Fig. 2F), as previously observed by TEM. While cells with fully formed septa were readily visible, we there was a smaller proportion of cells with incomplete septa compared to the TEM images, likely due to resolution limits of light microscopy, and the Nile Red staining only the membrane rather than the whole envelope. The 1771 compound reduced the proportion of cells with mature septa in both the wild type and MspA-deficient cells. However, the effect was more apparent in the absence of MspA where the proportion decreased from 30% to 21%, compared to the wild-type which showed a decrease from 15% to 12% (Fig. 2F). Autolysins are peptidoglycan hydrolases which support growth and division and mediate daughter cell separation after septum formation (Wang, Buist and van Dijl, 2022). Both WTA and LTA, and their D-alanylation, have been implicated in controlling the activity and localisation of autolysins (Schlag *et al*., 2010; Biswas *et al*., 2012; Zoll *et al*., 2012). Therefore, we employed an autolysis assay to test whether autolytic activity was altered in the absence of MspA, and if the increased amount of LTA could be responsible for it. Detergent-induced autolysis of the MspA-deficient cells was indeed significantly reduced compared to the wild type (Fig. 2F), suggesting dysregulation of the autolytic enzyme activity. Crucially, addition of compound 1771 to the medium restored the autolytic activity to the MspA-deficient cells (Fig.2F), while it did not significantly reduce autolysis of the wild-type cells. This suggests that the excess LTA causes dysregulation of the autolytic enzymes activity which, in turn, is likely responsible for the observed delay in autolytic splitting of daughter cells.

### The MspA protein interacts with the LTA synthesis enzymes

We have established that MspA-deficiency triggers overproduction of LTA which, in turn, deregulates autolytic activity important for daughter cell separation, a process that ultimately leads to an increased cell size. To understand this in greater detail, we next investigated how MspA is affecting LTA synthesis. Structural predictions suggest that MspA is a membrane protein and we verified this by fusing the *mspA* coding sequence with mCherry and performing fluorescence microscopy. This analysis revealed that MspA indeed localises to the membrane in a uniform pattern (Fig. 3A). The slightly more intense signal observed at the septum in dividing cells is expected due the presence of two membrane layers at that site. Initially we hypothesised that MspA could affect LTA synthesis by contributing to the stability of functional membrane microdomains (FMMs). These are domains in bacterial membranes similar to the eukaryotic lipid rafts, in *S. aureus* formed by staphyloxanthin and the scaffold protein flotillin (FloA) (López and Kolter, 2010; García-Fernández *et al*., 2017). FloA acts as a chaperone for the assembly of several protein complexes and as a consequence disruption of FMMs affects multiple phenotypes such as resistance to beta-lactams (García-Fernández *et al*., 2017), virulence (Koch *et al*., 2017) and function of the type VII secretion system (Mielich-Süss *et al*., 2017). Given that the deletion of *mspA* also has pleiotropic effects and that LtaS has been shown to be part of the FMMs protein cargo (García-Fernández *et al*., 2017), we decided to test whether MspA contributed to FMMs organisation by performing a bacterial two-hybrid assay between MspA and FloA. The MspA protein is predicted to have four transmembrane domains, with both the N- and C-termini exposed in the extracellular space (Fig. 3B). To ensure that the T18 and T25 tags located intracellularly such that interactions with another tagged protein can catalyse the necessary conversion of ATP to cAMP (Battesti and Bouveret, 2012), we cloned the protein’s last three transmembrane domains in the constructs where MspA was fused with these tags at the N-terminus and the first three transmembrane domains in the constructs where MspA was fused with the tags at the C-terminus (Fig. 3B). We did not observe any interactions between MspA and FloA using this approach (Fig. S5), suggesting that MspA doesn’t contribute to FMMs formation which is further supported by the difference in distribution of these protein across the membrane with MspA being evenly distributed (Fig. 3A), whereas FloA has a punctuated membrane localisation pattern (López and Kolter, 2010; García-Fernández *et al*., 2017). We next hypothesised that MspA could interact directly with the LTA synthesis enzymes to affect LTA production, in particular with LtaA, UgtP and LtaS which also localise to the membrane. To test this we combined the MspA bacterial two-hybrid constructs with the LtaA, UgtP and LtaS ones, except for the constructs where LtaS was tagged at the C-terminus, as this is known to be located extracellularly. MspA was found to interact with all three proteins (Fig. 3C, Fig. S6), with the colony colour intensity suggesting a stronger interaction with LtaA and LtaS.

**Figure 3.**
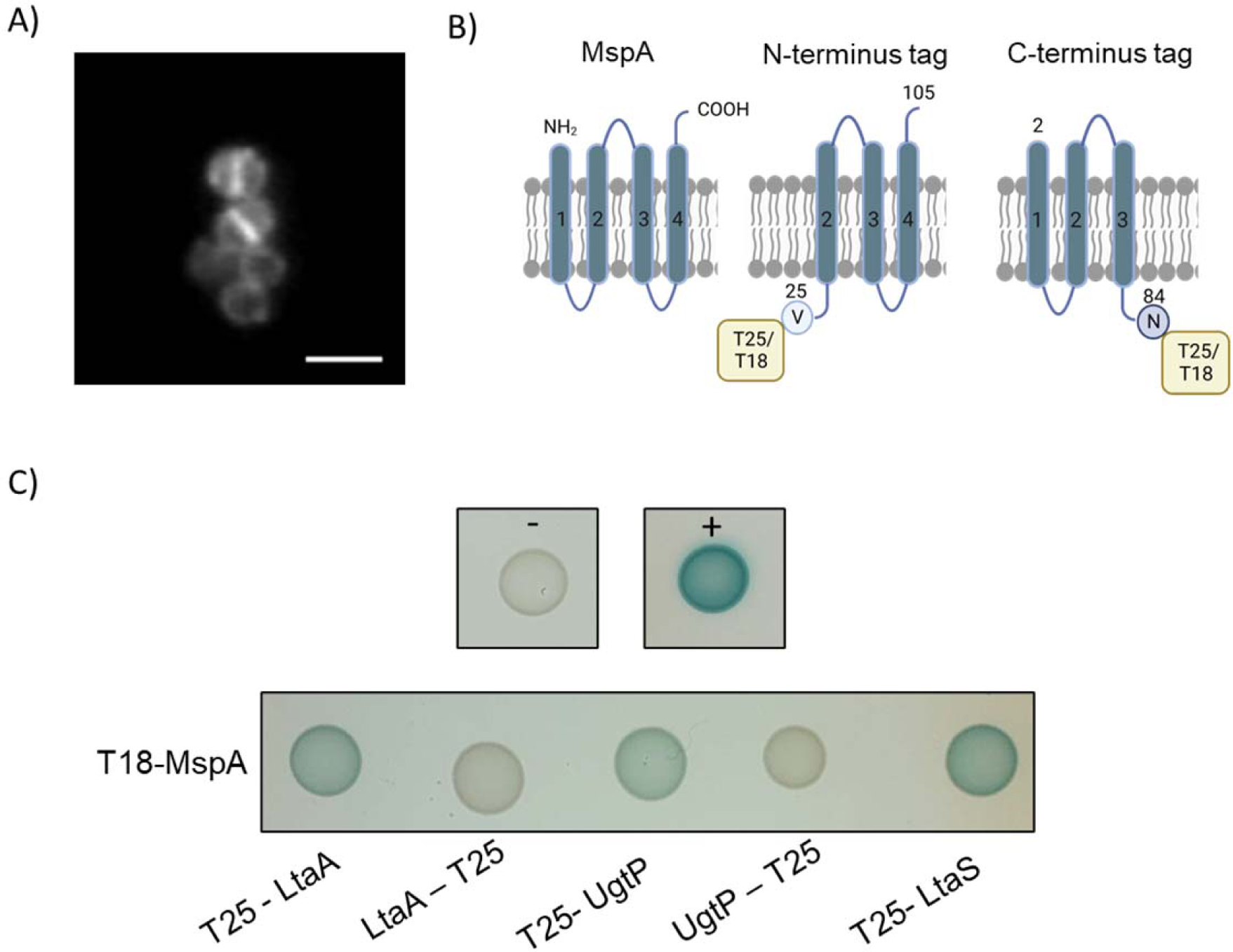
MspA interacts with the LTA synthesis enzymes. A) Widefield fluorescent microscopy of SH1000 pOS1 MspA-mCherry showing localisation of MspA at the cell membrane. Scale bar: 2 µm B) Diagram of the MspA protein and of its truncated versions which were cloned in the bacterial two-hybrid plasmids to maintain the T18 or T25 tag intracellularly. Created with BioRender. C) Bacterial two-hybrid analysis between MspA tagged with the T18 subunit at the C-terminus and LtaA, UgtP and LtaS tagged with the T25 subunit at the N-or C –terminus, as indicated. LtaS was tested only with the tag at the N-terminus as the C-terminus localizes extracellularly. MspA interacts with LtaA, UgtP and LtaS tagged at the N-terminus. Representative of three biological replicates. pKT25 and pUT18 plasmids were co-transformed as negative control (-) and pKT25 – zip and pUT18C – zip as a positive control (+).

### MspA interferes with the interaction between LtaA and LtaS

Having showed that MspA interacts with UgtP, LtaA and LtaS, we hypothesised that MspA could affect the stability of the LTA enzymatic complex. To test this, we performed a bacterial three-hybrid assay investigating whether MspA could alter the association between LtaA and LtaS specifically, as MspA had a stronger interaction with these two proteins. First, we confirmed with a bacterial two-hybrid the positive interaction between T25 - LtaS and LtaA, tagged with the T18 subunit both at the C-or N-terminus (Fig. 4A). We then complemented this set up with a third plasmid, pSEVA641, expressing *mspA* under the *lac* promoter. Expression of MspA supressed the interaction between T25 – LtaS and both LtaA-T18 and T18 – LtaA, compared to co-transformants carrying the empty pSEVA641 plasmid (Fig. 4A, Fig. S7). This suggests that MspA could have a role in preventing the formation of the LTA synthesis complex or in favouring its disassembly.

**Figure 4.**
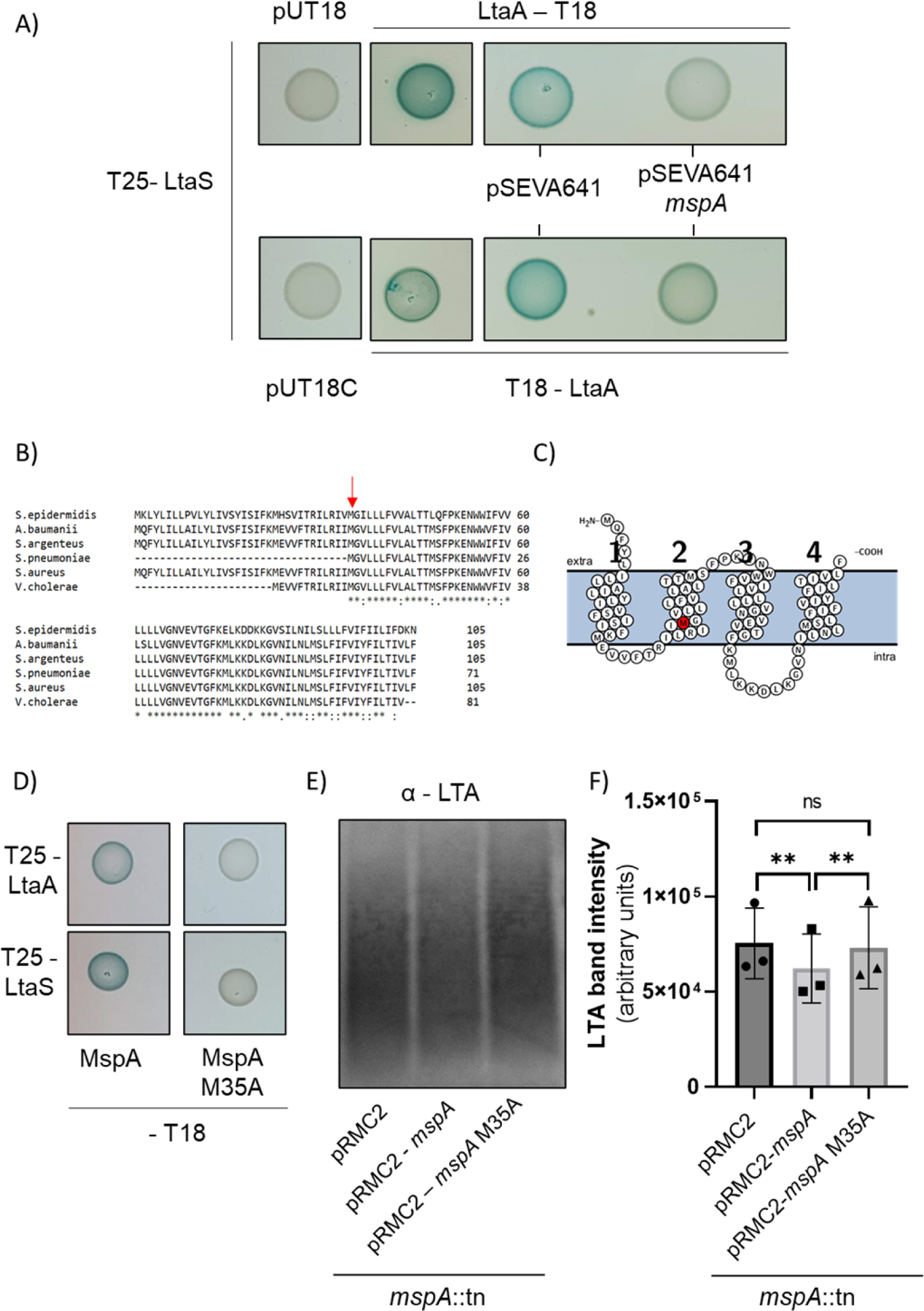
MspA affects LTA production by interfering with the interaction between LtaA and LtaS. A) Bacterial three-hybrid analysis testing interaction between T25-LtaS and the empty plasmids pUT18 and pUT18C as a negative control, between T25 – LtaS with LtaA tagged with T18 at the N-or C-terminus alone and T25-LtaS with T18-LtaA or LtaA-T18 and a third plasmid, either the empty pSEVA641 or pSEVA641 expressing *mspA* under the *lac* promoter. Representative of 3 biological replicates, see figure S7. B) Alignment of MspA protein sequence from *S. aureus* with homologues found in *Staphylococcus epidermidis*, *Acinetobacter baumaniii*, *Staphylococcus argenteus*, *Streptococcus pneumoniae* and *Vibrio cholerae*. The conserved residue methionine 35 is highlighted with a red arrow. C) Diagram of the MspA protein showing the localisation of its amino acids, extracellularly (extra), transmembrane or intracellularly (intra) as predicted by Protter (Omasits *et al*., 2014). Highlighted in red in the second transmembrane domain with its key residue methionine 35. D) Bacterial two-hybrid assay between LtaA or LtaS and wild-type MspA (MspA) or MspA with methionine residue 35 mutated to alanine (MspA M35A). E) Western blot with anti-LTA antibodies on samples from SH1000 *mspA*::tn complemented with empty pRMC2, pRMC2-*mspA* with a wild-type copy of *mspA* and pRMC2-*mspA* M35A with a copy of *mspA* with methionine residue 35 mutated to alanine. F) Quantification of the intensity of LTA western blot bands performed with ImageJ (*n*= 3, one-way ANOVA with Tukey *post-hoc* test, pRMC2 vs pRMC2-*mspA*, **, P = 0.002; pRMC2-*mspA* vs pRMC2-*mspA* 35A, **, P = 0.005; pRMC2 vs pRMC2-*mspA* M35A,ns, P = 0.4).

To further verify that native MspA can interfere with LtaA and LtaS interactions we identified and mutated a key residue. MspA is highly conserved across *S. aureus* strains but it is also present in a number of distally related species. Protein sequence alignment of MspA from *S. aureus* with the homologues of five other species (Fig. 4B) highlighted multiple conserved amino acids, likely important for the protein activity. The mutation of the first of these residues, methionine 35, situated in the second transmembrane domain (Fig. 4C), to alanine indeed abolished the interaction between MspA and LtaA and LtaS (Fig. 4D, Fig. S8), suggesting that this residue was essential for MspA’s interaction with the LTA enzymes. We then tested whether mutated MspA (MspA M35A) could rescue the increase in LTA observed in the absence of MspA. While expression of wild-type MspA decreased the amount of LTA, the strain expressing MspA M35A exhibited LTA-content comparable to the empty plasmid control (Fig. 4E, 4F, Fig. S9). These observations show that the interaction of MspA with the MspA enzymes is critical for MspA’s ability to maintain LTA homeostasis. They also provide evidence that MspA acts by interfering with the interactions between the LTA enzymes, thereby reducing LTA biosynthesis.

## DISCUSSION

We have previously shown that MspA is critical to the virulence of *S. aureus* due to its effects on toxin production, membrane stability, staphyloxanthin biosynthesis, iron homeostasis and susceptibility to components of the innate immune system (Duggan *et al*., 2020). In this study, we decipher the molecular basis of this cellular role. We show that MspA directly interacts with enzymes of the LTA synthetic pathway in a competitive manner. This interaction contributes to LTA homeostasis which, in the absence of MspA, leads to an increase in cell size likely due to a delay autolytic daughter-cell separation. A graphical summary of this process is provided (Fig. 5).

**Figure 5.**
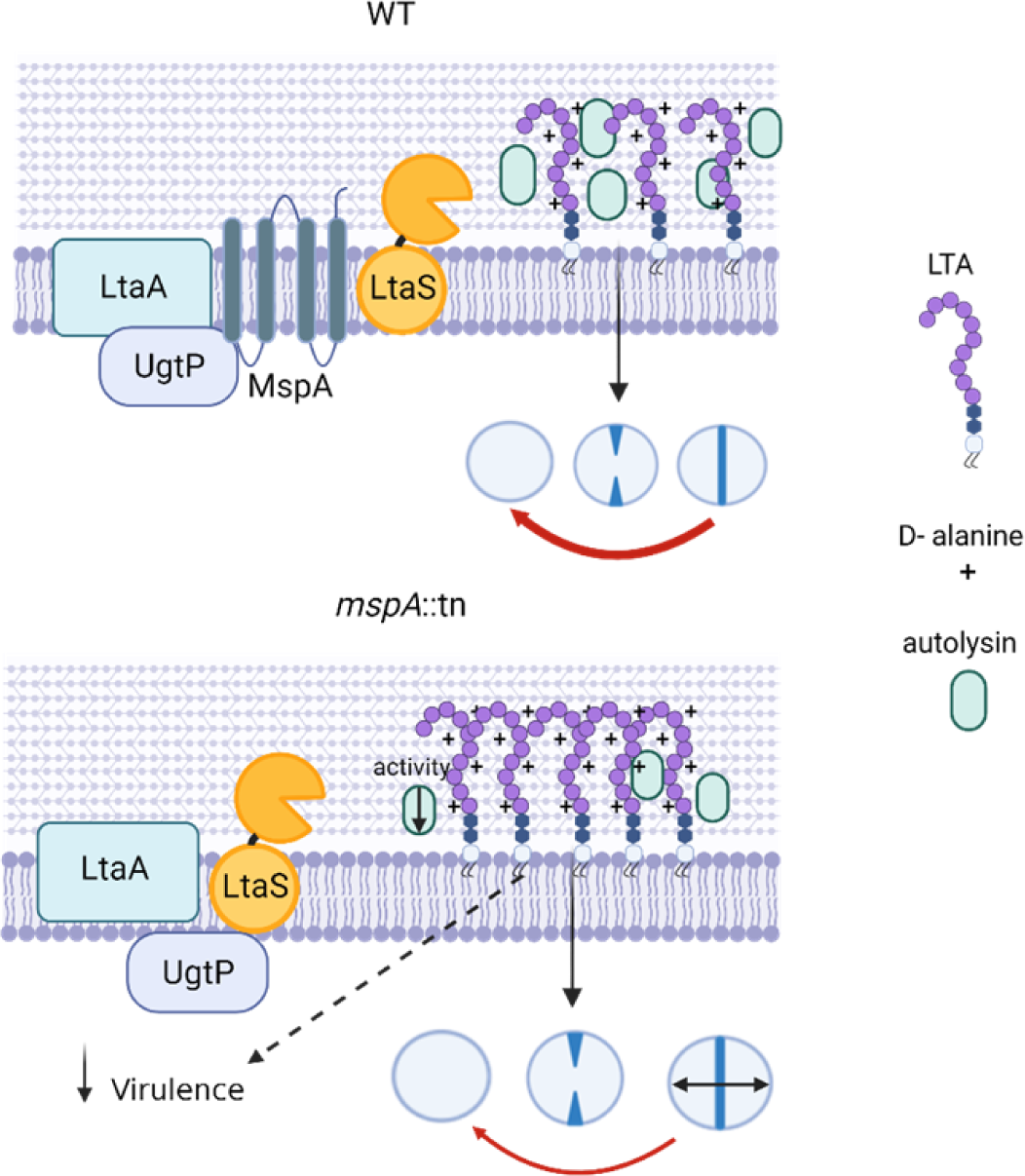
MspA negatively regulates LTA production by competitively interacting with the LTA enzymes. MspA maintains normal cell cycling and cell size by regulating LTA synthesis and, indirectly, autolysis activity. The LTA defects observed in the absence of MspA could also underpin the reduced virulence.

Autolytic enzymes affect both cell morphology and the cell cycle progression. In *S. aureus*, deletion of Atl and Sle1 autolysins leads to elongation of the cell cycle (Monteiro *et al*., 2015) while Sle1-inactivation also causes a delay in cell daughter separation, resulting in an increase in cell size similarity to what is observed in the absence of MspA (Monteiro *et al*., 2015; Thalsø-Madsen *et al*., 2020; Veiga *et al*., 2023). An *atl* deletion mutant, in contrast, exhibits initiation of new septa before the previous round of cell division is completed, resulting in disturbed cell separation and cell clumping (Biswas *et al*., 2006; Chan *et al*., 2016; Nega *et al*., 2020). Similar formation of premature septa was also observed in a small number of cells deficient for MspA. In contrast to *sle1*, deletion of *atl* does not alter the cell size (Takahashi *et al*., 2002). However, upregulation of secondary hydrolases in upon absence of Alt has been suggested to compensate for defects caused by *atl* deletion (Pasztor *et al*., 2010; Hirschhausen *et al*., 2012). LTA is known to regulate the autolytic process although the detailed mechanisms are not well understood. LTA has previously been shown to bind the Atl amidase repeats, likely targeting Atl to the septum and modulating its activity, as increasing concentration of LTA decrease the peptidoglycan amidase activity (Zoll *et al*., 2012). Accordingly, other mutants with altered LTA synthesis display defects in autolysis. *ugtP* deletion mutants were shown to have reduced autolytic activity (Fedtke *et al*., 2007) and Δ*ltaS* strains with suppressor mutations were reported to have decreased abundance of autolysins in the cell wall (Oku *et al*., 2009; Corrigan *et al*., 2011). Lack of UgtP and LtaA leads to longer LTA polymers which, similar to lack of MspA, lead to an increased cell size due to defects in cell cycle progression. In the case of UgtP and LtaA, however, the septum formation rather daughter cell separation is delayed (Hesser, Matano, *et al*., 2020). This shows that LTA abundance and length can affect the cell cycle at different stages.

Here, we provide evidence that MspA contributes to maintaining physiological levels of LTA in the cell wall. There are two other known mechanisms that negatively regulate LTA production, confirming the importance of controlling and balancing the polymer synthesis: (i) post-translational inactivation of the LtaS enzyme by proteolytic cleavage of its extracellular domain by the SpsB signal peptidase (Wörmann *et al*., 2011) and (ii) recently discovered transcriptional downregulation *ltaS*-expression in the stationary phase of by the essential two-component system WalKR (Sharkey *et al*., 2023).

While LtaS localises mainly at the septum, LtaA and UgtP are distributed more homogenously along the membrane, suggesting that glycolipid anchor synthesis could happen independently from LTA synthesis (Reichmann *et al*., 2014). MspA’s cellular localisation resembles that of LtaA and UgtP. Therefore, MspA has the opportunity to interact with all three proteins at the septum where LTA chain synthesis takes place to regulate the polymer abundance, as well as associate solely with LtaA and UgtP throughout the membrane. While further work is needed to elucidate the mechanistic details, we suggest that interference with the LTA biosynthetic enzymes’ interactions by MspA could prevent LTA biosynthetic enzyme complex formation, or favour its disassembly. In the absence of MspA, the complex would be stabilised, leading to an increase in LTA production. Additionally, it is also possible that MspA could affect the processivity of the LTA enzymes.

The synthesis and turnover of cell envelope components are known to impact virulence and pathogenicity of *S. aureus*. For instance, the membrane lipid lysyl-phosphatidylglycerol (LPG), LTA (Zheng *et al*., 2021) and WTA (Brignoli *et al*., 2022) control sorting and secretion of toxins, and deletion of the scaffold protein FloA causes decrease in virulence as it is needed to stabilise RNase Y, which degrades small RNAs downregulating toxins (Koch *et al*., 2017). Additionally, LTA has been shown to impact virulence, as *ltaA* and *ugtP* deletion mutants have attenuated pathogenicity (Gründling and Schneewind, 2007a; Sheen *et al*., 2010). We hypothesise that increased abundance of LTA in the absence of MspA causes major structural changes to the membrane that has pleiotropic effects on virulence. In conclusion, we have shown that MspA is a membrane protein that competitively interferes with LTA synthetic enzymes, leading to a dysregulation of LTA biosynthesis with major pleiotropic consequences for the ability of the bacteria to cause disease. This work therefore uncovers a new link between envelope synthesis and *S. aureus* virulence, and suggests that MspA could represent a promising target for future therapeutic development.

## METHODS

### Strains and culture conditions

*Staphylococcus aureus* strains were grown in TSB at 37°C at 180 rpm shaking unless specified otherwise. When appropriate, chloramphenicol (10 µg/ml) and anhydrous tetracycline (200 ng/ml) (Thermo Fisher Scientific) were added. *E. coli* strains were grown in LB, Mach1 and DH5α cloning strains at 37°C at 180 rpm shaking, and BTH101 at 30 °C at 180 rpm shaking. When appropriate, ampicillin (100 µg/ml), kanamycin (30 µg/ml) or gentamycin (10 µg/ml) were added.

### Genetic manipulations

Phusion polymerase (Thermo Fisher Scientific) was used for PCR amplification. New England Biolabs enzymes were used for restriction digestion and for ligation.

### Electron microscopy

Strains were cultured at 37°C in 5 ml TSB in a 50 ml falcon tube for 18 hours. An 800 µl volume was removed and pelleted. Cells were fixed by resuspending pellets in 2.5% glutaraldehyde in cacodylate buffer (pH 7.3) and stored at 4°C until further processing. Fixed pellets were resuspended in BSA/glutaraldehyde gel at 10-20°C and once again pelleted. Pellets were postfixed with osmium ferrocyanide/osmium tetroxide in cacodylate buffer, stained with 2% uranyl acetate and Walton’s lead aspartate. Finally, samples were subject to ethanol dehydration, then infiltration with propylene oxide and EPON resin mix. Embedded blocks were polymerised for 48 hours at 60°C. Multiple 70 nm sections were cut on a Leica UC7 ultramicrotome and imaged using a FEI Tecnai T12 microscope at the Wolfson Bioimaging Facility at the University of Bristol.

### Analysis of LTA via Western Blot

Strains were grown overnight and 1 ml of culture was transferred to a 2 ml Lysing matrix B tube (MP Biomedicals) and bead-beaten for 1 minute at 5 m/s twice. Beads were settled via centrifugation at 2000 rpm in a tabletop centrifuge and 500 µl of supernatant centrifuged at 13,000 rpm for 15 minutes. The supernatant was discarded and the pellet was resuspended in a 1:1 solution of 100 mM Tris-HCl pH 7.4 with EDTA-free protease inhibitor cocktail (Roche, 1 tablet in 10ml of buffer) and 4x NuPage LDS sample buffer (Invitrogen) to normalise to the equivalent of OD_600_ 6 in 100 µl. The resuspended pellets were boiled for 20 minutes and centrifuged at 13,000 rpm for 5 minutes. The supernatant was harvested and stored at −20 °C. Samples were heated at 70°C for 10 minutes and loaded on FastGene PAGE Gel, 4-12% (GeneFlow). Volumes loaded were optimised between 7 to 12 µl to obtain even protein loading. Gels were run in MES buffer (NuPage) and stained with Quick Coomassie Stain (NeoBiotech) overnight or transferred to methanol activated PVDF 0.2 µm membrane (Cytiva) in a semi-dry transfer at 25 V for 30 minutes with transfer buffer (600 mM Tris, 600 mM glycine, 280 mM Tricine, 0.05% SDS and 2.5 mM EDTA). Membranes were blocked with 3% BSA in PBS, stained with 1:1000 α-LTA mAb 55 (Hycult BioTech) and with 1:10,000 anti-mouse IgG Peroxidase HRP (Sigma Aldrich). 50 µg/ml of IgG from human serum (Merck) in PBS were added to all blocking and antibody incubations. Membranes were developed with Metal Enhanced DAB Substrate Kit (Thermo Fisher Scientific). Blot images were taken with Genesys acquisition system (Syngene) and band intensity was quantified with ImageJ.

### 1771 MIC

Strains were grown overnight in overnight and diluted to OD_600_ of 0.05 in 50 ml Mueller-Hinton broth and grown to OD_600_ of ∼ 0.5. Cultures were diluted to OD_600_ of 0.005 in 10 ml of Mueller-Hinton. 100 µl of culture in triplicate were added to 100 µl of Mueller-Hinton broth with 2-fold dilutions of 1771 compound. Plates were incubated at 37°C for 24 hours and OD_600_ recorded with SPECTROstar Nano plate reader.

### Fluorescence microscopy

For imaging of membrane-stained *S. aureus*, 1 ml of bacteria grown overnight with or without 2 µg/ml 1771 compound were washed 3 times in PBS and resuspended in 1ml PBS with 10 µg/ml Nile Red. Samples were incubated at 37 °C in the dark for 5 minutes, washed 3 times in PBS, resuspended in 4% paraformaldehyde in PBS, incubated at room temperature for 15-30 minutes, and stored at 4°C in the dark. For image acquisition, 2 µl of sample were mounted on a slide with 1.2 % agarose and covered with a cover slip. Images were taken with a with Leica SP8 AOBS confocal laser scanning microscope attached to a Leica DM I8 inverted epifluorescence microscope. Images were analysed with ImageJ Fiji (Schindelin *et al*., 2012).

For the analysis of MspA localisation, SH1000 cells carrying the expression plasmid pOS1 *mspA-mCherry* were grown overnight in TSB, followed by 1:100 dilution in fresh medium and incubation at 30 °C upon shaking. 0.5 µl of mid-logarithmic growth phase culture was transferred to 1.2% agarose-H_2_0 slides followed by microscopy. The imaging was carried out with Nikon Eclipse Ti-E microscope equipped with Nikon CFI APO TIRF × 100/1.49 objective, Cobolt Jive 100 561 nm solid-state laser light source and Andor iXon Ultra 897 EMCCD camera. Images were acquired with Nikon NIS Elements AR 5.11, and processed with Fiji using PureDenoise algorithm (Luisier *et al*., 2010; Schindelin *et al*., 2012).

### Autolysis assay

Strain were grown overnight and diluted to OD_600_ of 0.05 in 5 ml of TSB medium in the presence and absence of 2 µg/ml of 1771, followed by incubation until an OD_600_ of ∼ 0.3 - 0.5 and a single wash in ice-cold water. The samples were normalized to OD_600_ of 1 in water supplemented with 0.1% Triton X-100. 200 µl of culture was transferred to a flat bottom 96 well plate in triplicate and OD_600_ was monitored over 6 hours every 30 minutes with 500 rpm shaking before each reading with SPECTROstar Nano plate reader.

### Bacterial two-hybrid

Genes were amplified from *S. aureus* JE2 genomic DNA with primers listed in Table S3 and mutated *mspA* sequences were amplified from the corresponding pRMC2-*mspA* plasmids. Amplification products and pKT25, pKNT25, pUT18 and pUT18C plasmids were digested with *BamH*I and *Kpn*I enzymes, ligated and 2 µL of ligation mixture were transformed into Mach1 or DH5α competent cells and screened for insertion with primers BTH_F and pKT25/NT25_R or pUT18/18C_R. One positive clone per construct was grown overnight for plasmid extraction. Correct insertion was checked by amplifying the construct with BTH_F_primer and reverse primer matching the insert, followed by sequencing. Pairwise combinations of plasmids were co-transformed in BTH101 *E. coli* cells. Three colonies per co-transformation were grown overnight with appropriate antibiotics and 1 mM of IPTG and 10 µl per replicate was dotted on LB containing antibiotics, 50 µg/ml of X-gal and 1 mM IPTG. Plates were incubated at 30 °C for 24 hours.

### Bacterial three-hybrid

The *lac* promoter and the *mspA* gene were amplified with primers listed in Table S3 from pKT25 plasmid and JE2 genomic DNA respectively. The two fragments were fused with PCR with the external primers. The fusion product and plasmid pSEVA641-*floA* were cut with *EcoR*I and *Hind*III restriction enzymes, ligated and 2 µl of ligation mixture were transformed into Mach1 cells, plated on 10 µg/ml gentamycin. Colonies were screened for insertion with primers pSEVA MCS reverse and Lac promoter EcoRI F and a positive clone was grown overnight for plasmid extraction and sequencing. Empty pSEVA641 plasmid was obtained by cutting pSEVA641-*floA* at the *Not*I site, re-ligated and transformed in Mach1. One colony per transformation was grown overnight for plasmid extraction and sequencing. Three-plasmid combinations were co-transformed in BTH101 *E. coli* cells. Three colonies per co-transformation were grown overnight with appropriate antibiotics and 1 mM IPTG. 10µl per replicate was dotted on LB containing antibiotics, 50 µg/ml of X-gal and 1 mM IPTG. Plates were incubated at 30 °C for 24 hours.

### MspA site directed mutagenesis

Single site mutation M35A was generated in the *mspA* gene amplifying the pRMC2-*mspA* plasmid with the primers listed in Table S3. PCR products were digested with DpnI and transformed into Mach1 (Invitrogen). Three colonies per transformation were grown and used for plasmid purification. Single site mutation was checked via sequencing.

### Statistical analyses

Statistical analyses indicated in the figure legends were performed using GraphPad Prism 9.2.

## Acknowledgments

We thank Angelika Gründling for providing the bacterial two-hybrid plasmids, protocols for lipoteichoic acid detection and for helpful discussions to guide the project. We thank Maisem Laabei for providing us with the compound 1771 and Daniel López for providing us with the pSEVA641-*floA* plasmid. We also acknowledge Lorna Hodgson and Katy Jepson from the Wolfson Bioimaging Facility for training and guidance on the use of the confocal microscope.

## Supplementary Figures

**Figure S1.**
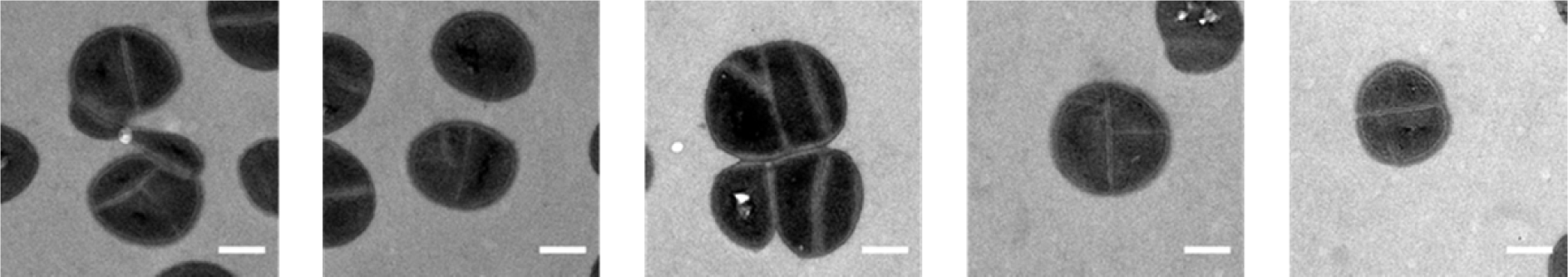
TEM images of SH1000 *MspA* deficient mutant cells with irregular septa. A small number of SH1000 *mspA*::tn cells have multiple septa for cells and/or septa that are not perpendicular to the previous septal plane. These defects were not observed in SH1000 wild type. Scale bar: 0.5 µm.

**Figure S2.**
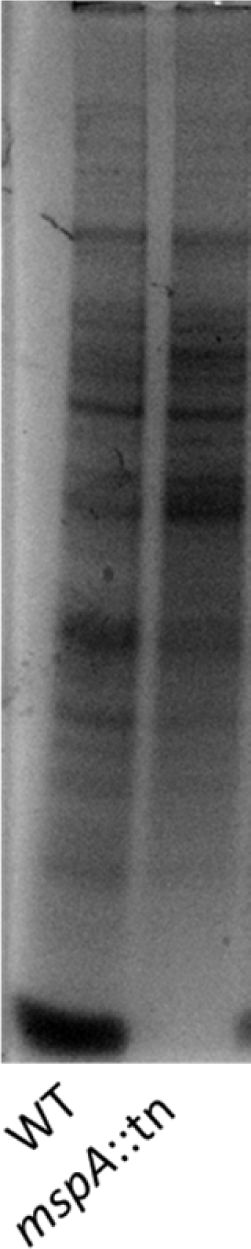
Coomassie stain of SDS-PAGE for protein loading control. Coomassie stain of SDS-PAGE with samples of SH1000 and *mspA* mutant showing total protein loading control for LTA western blot in Fig. 2A.

**Figure S3.**
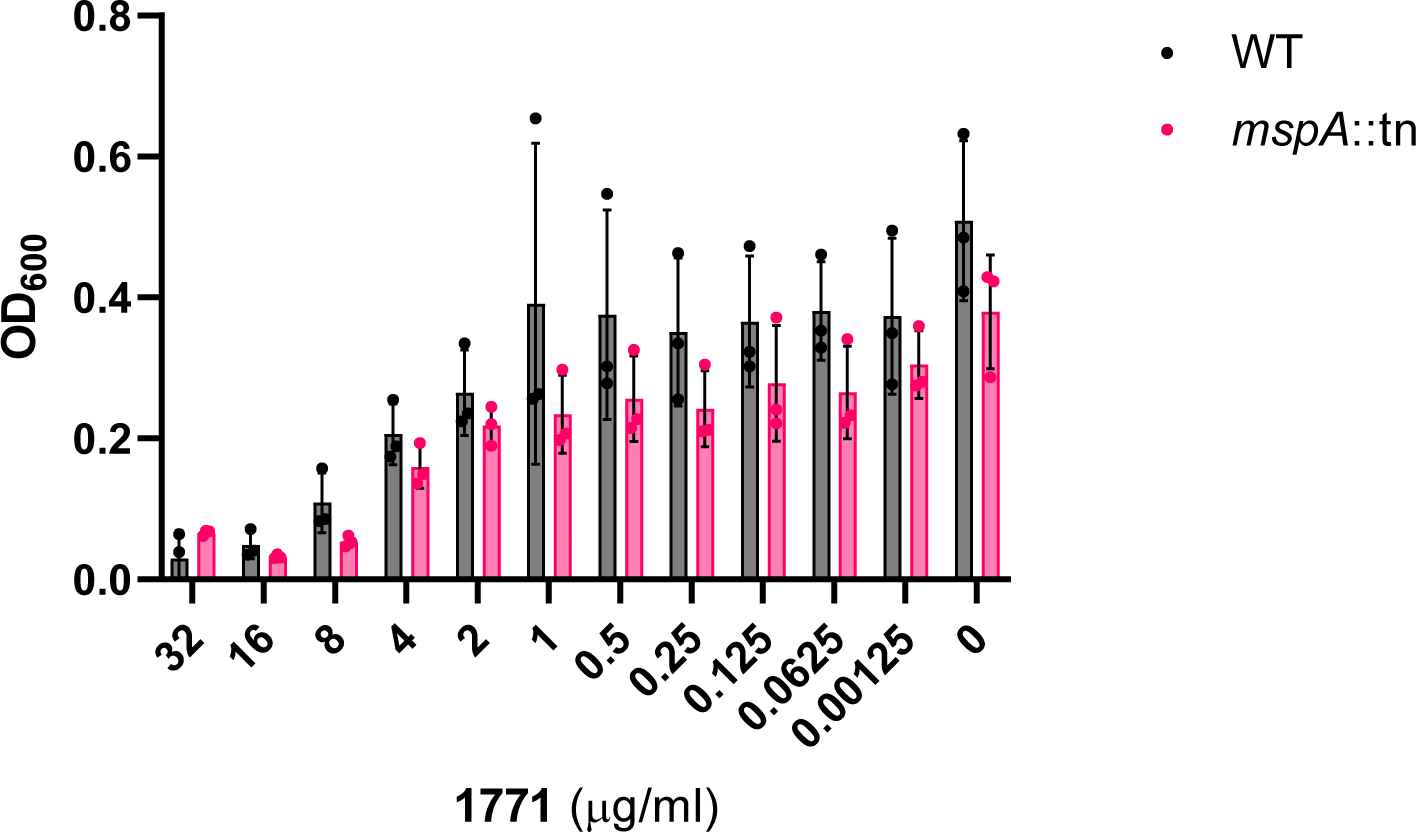
Minimum inhibitory concentration assay to test wild type and *mspA* mutant sensitivity to the 1771 compound. Each dot on the graph represents an individual biological replicate, bars represent ±SD.

**Figure S4.**
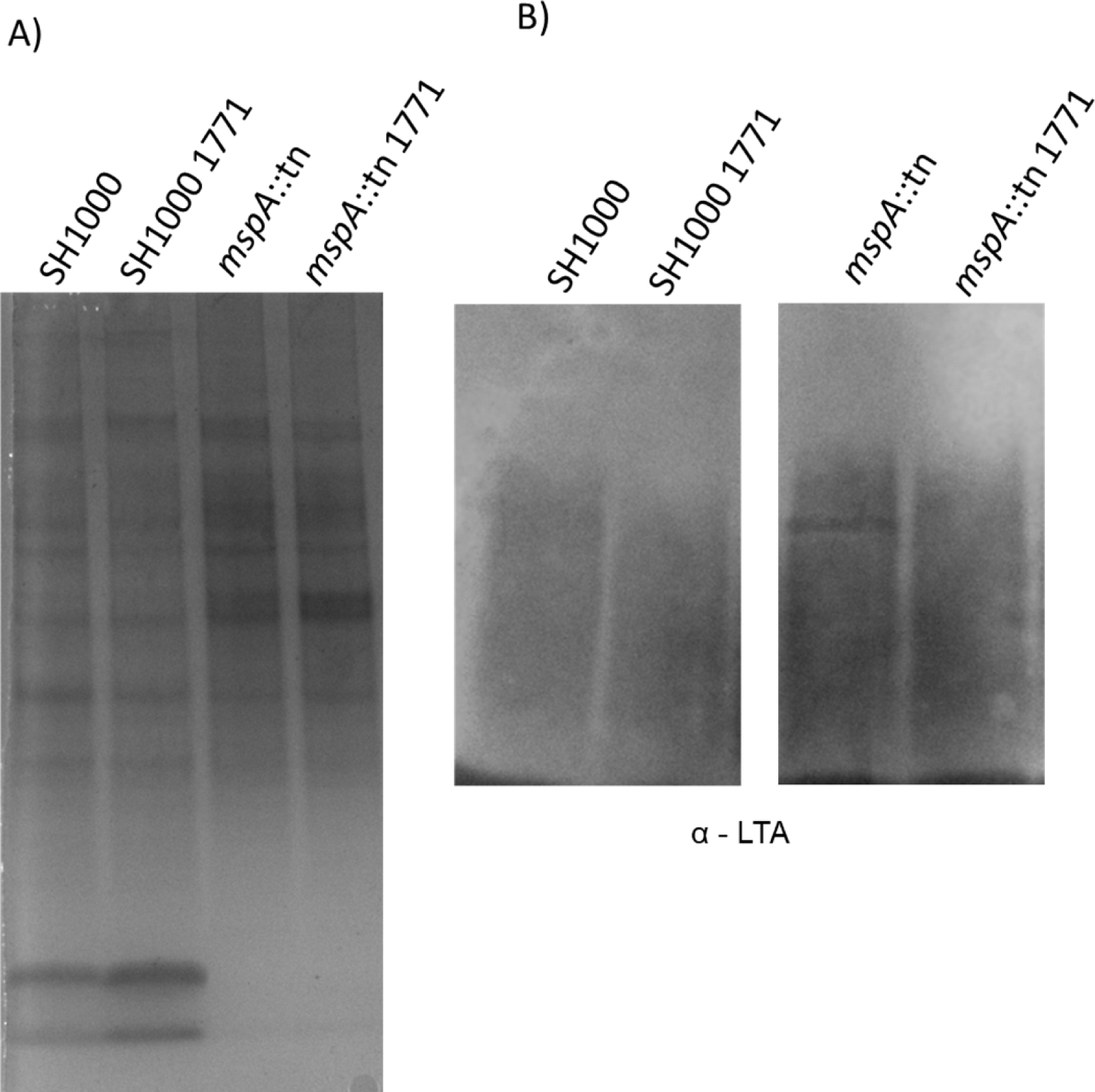
LTA production in wild type and *mspA* mutant grown with and without 1771 compound. Coomassie stain of SDS-PAGE (A) and Western blot with anti – LTA antibodies (B) of samples from SH1000 and SH1000 *mspA*::tn grown overnight with and without 1771 compound (2 µg/ml).

**Figure S5.**
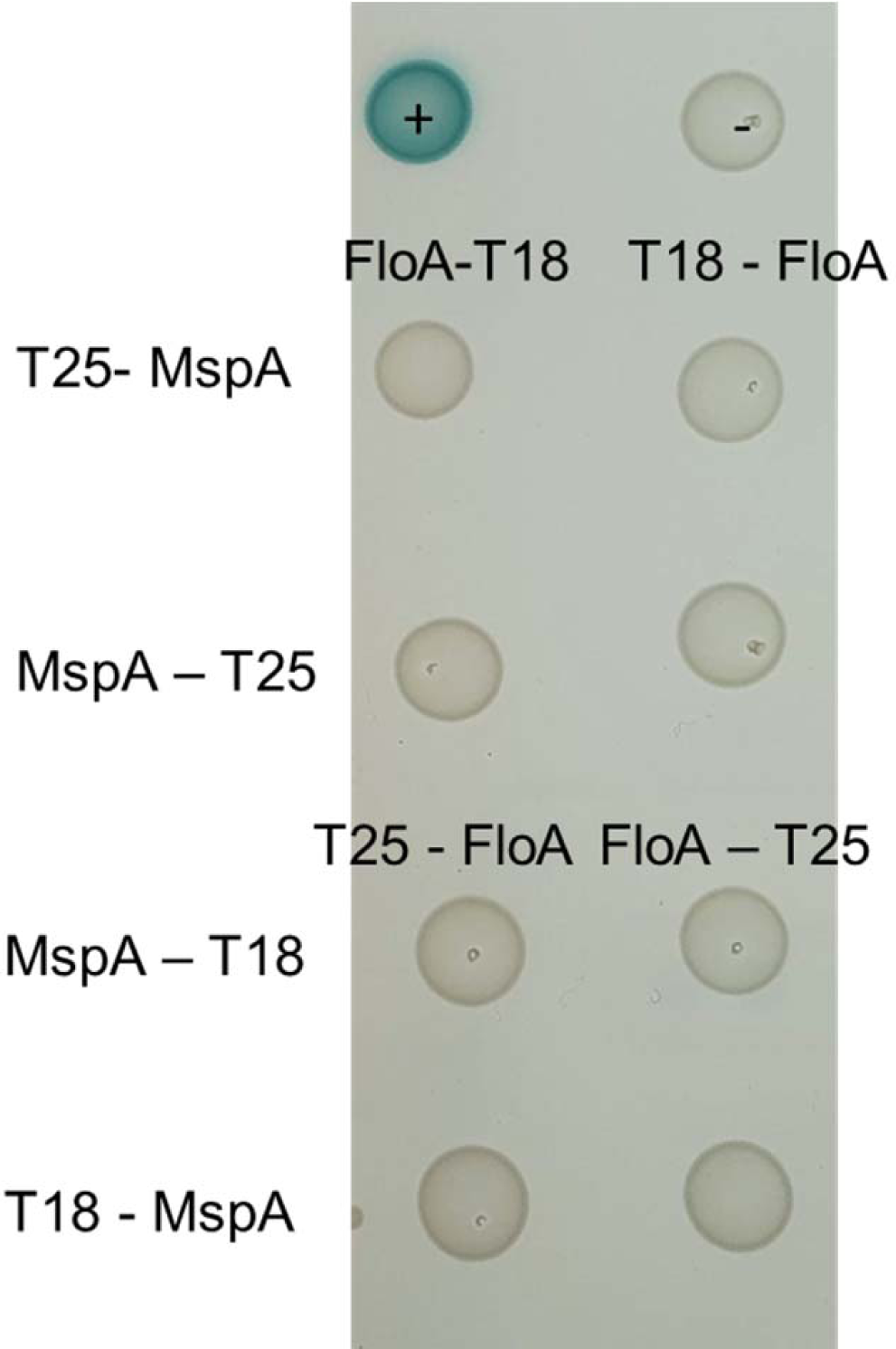
MspA does not interact with flotillin. Bacteria two-hybrid assay to test the interaction between MspA and flotillin (FloA). pKT25 and pUT18 plasmids were co-transformed as negative control (-) and pKT25 – zip and pUT18C – zip as a positive control (+). Representative of three biological replicates.

**Figure S6.**
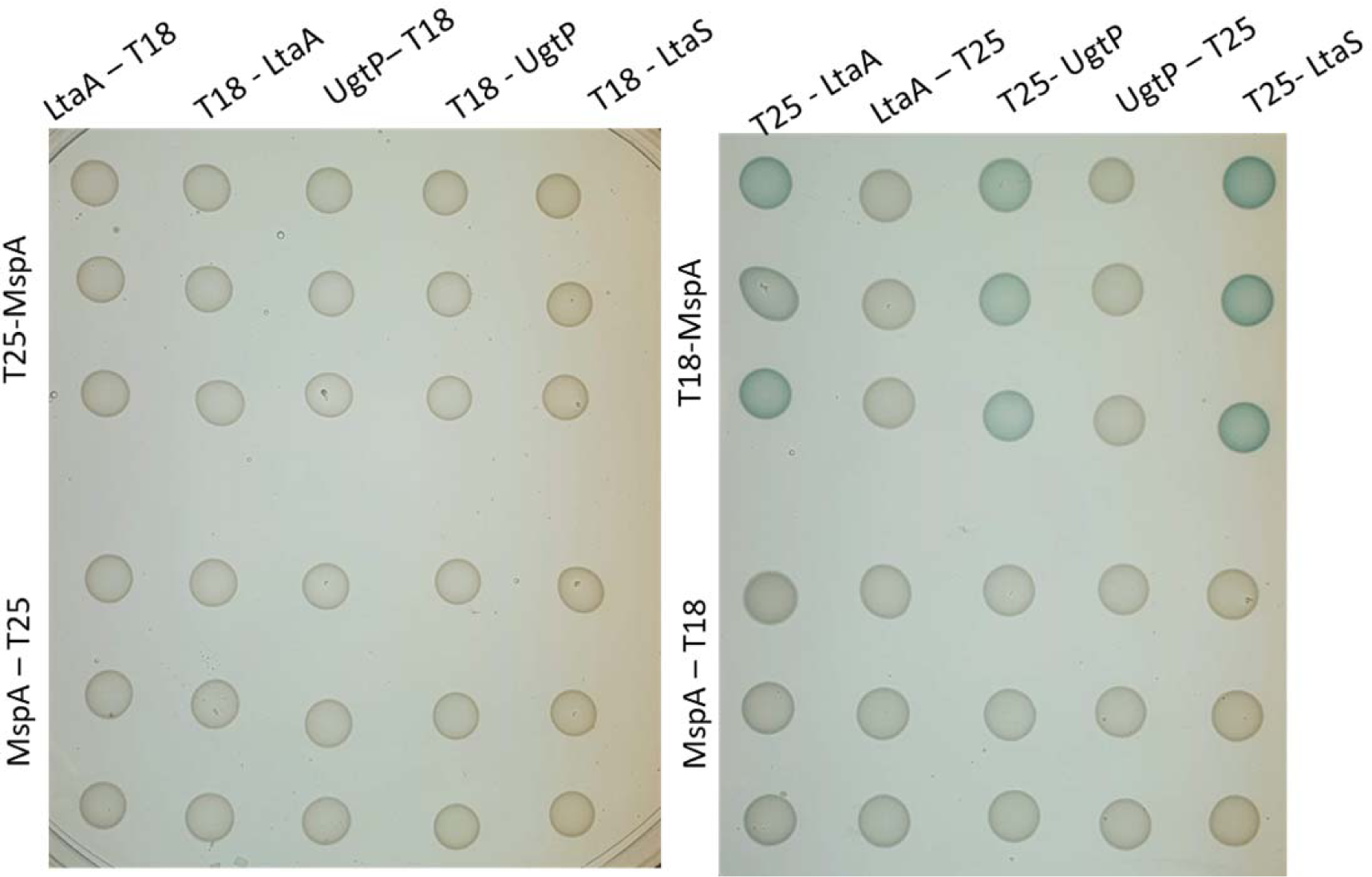
All bacterial two-hybrid interactions tested between MspA and UgtP, LtaA and LtaS. MspA (amino acids 25-105) tagged at the C-terminus with T18 interacts with UgtP, LtaA and LtaS when these are tagged at the N-terminus. LtaS was only tagged at the N-terminus as the C-terminus is known to localise extracellularly. Each dot for each combination of interactions tested represents a biological replicate.

**Figure S7.**
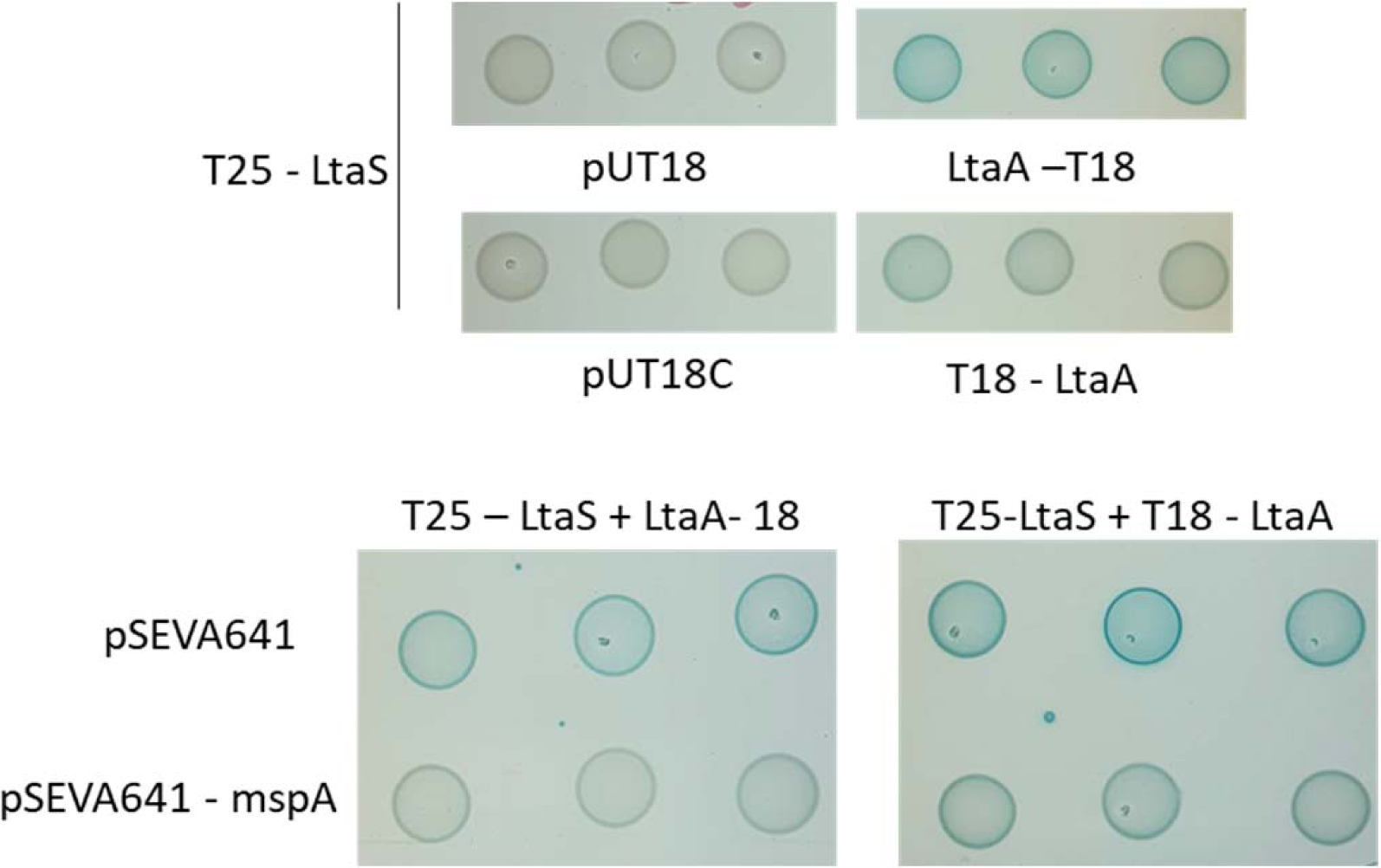
Bacterial three-hybrid testing the interaction between LtaS and LtaA in the presence and absence of MspA. Each dot for each combination of interactions tested represents a biological replicate.

**Figure S8.**
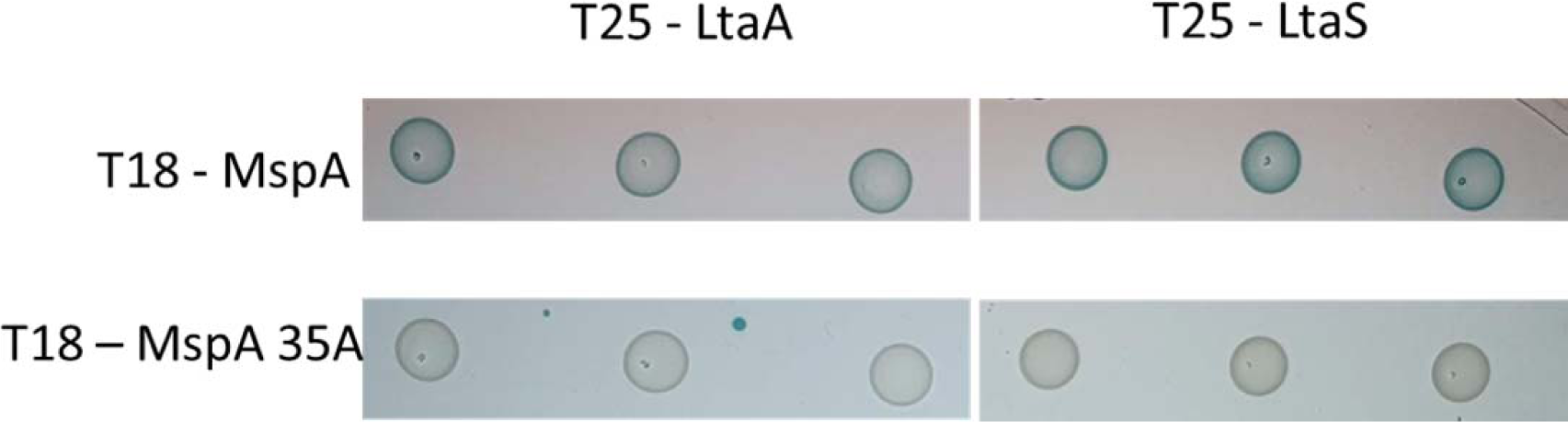
Bacterial two-hybrid testing the interaction between MspA or MspA 35A with LtaA and LtaS. Mutation of MspA residue methionine 35 to alanine (MspA 35A) disrupts the interaction between MspA and LtaA and LtaS. Each dot for each combination of interactions tested represents a biological replicate.

**Figure S9.**
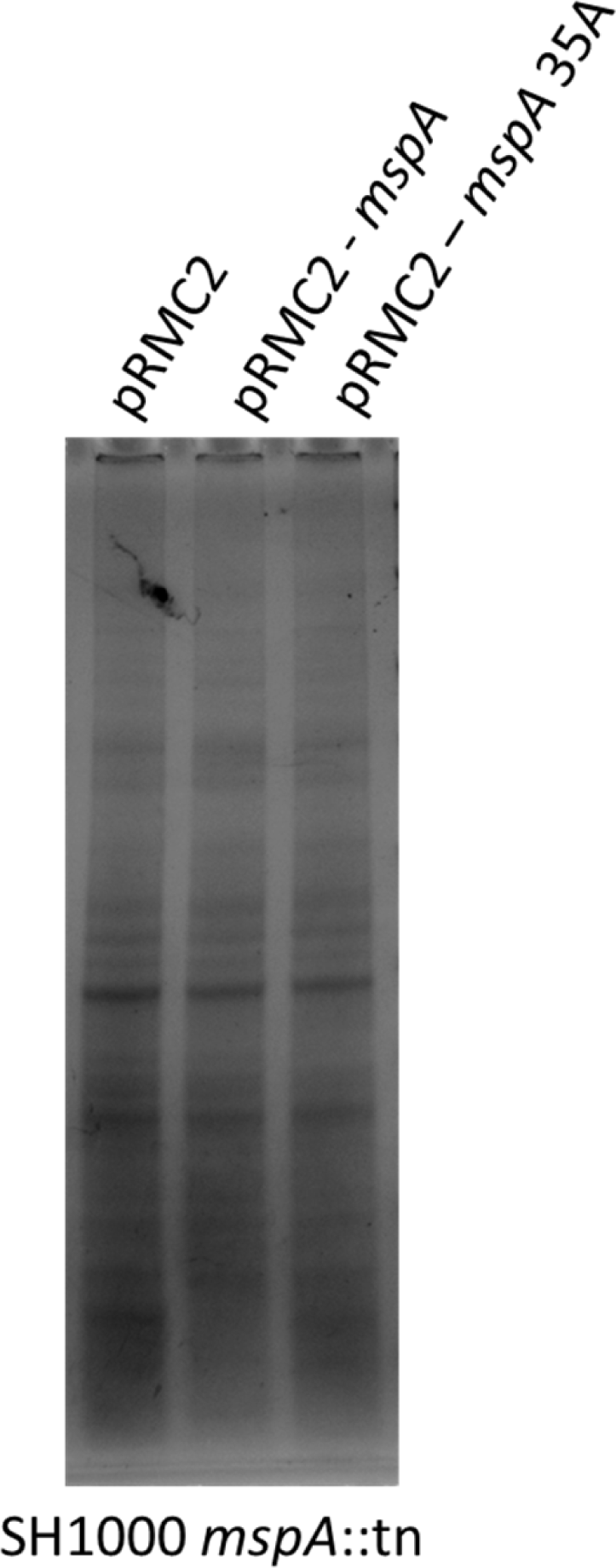
Protein loading control for LTA western blot. Coomassie stain of SDS-PAGE with samples of the indicated complemented SH1000 *mspA*::tn strains showing total protein loading control for LTA western blot in Fig. 4E.

## Supplementary tables

**Table S1.** Strain used in this study.

| Strain name | Description | Reference |
| --- | --- | --- |
| ***Staphylococcus aureus*** |  |  |
| SH1000 | Laboratory strain, 8325-4 with a repaired *rsbU* gene; SigB positive | (Horsburgh *et al.*, 2002) |
| SH1000 *mspA*::tn | *mspA* transposon mutant in SH1000 | (Duggan *et al.*, 2020) |
| SH1000 *mspA*::tn pRMC2-*mspA* | *mspA* transposon mutant in SH1000 complemented with *mspA* expressed on the pRMC2 plasmid | (Duggan *et al.*, 2020) |
| SH1000 pOS1 *mspA*-mCherry | SH1000 expressing *mspA* gene fused with mCherry on the pOS1 plasmid | This study |
| RN4220 | restriction-negative derivative of NCTC8325-4 | (Nair *et al.*, 2011) |
| ***Escherichia coli*** |  |  |
| Mach1 | Cloning strain | Laboratory collection |
| DH5α | Cloning strain | Laboratory collection |
| BTH101 | BACTH Δ*cya* strain; StrepR ANG1309 | (Karimova, Dautin and Ladant, 2005) |

**Table S2.**
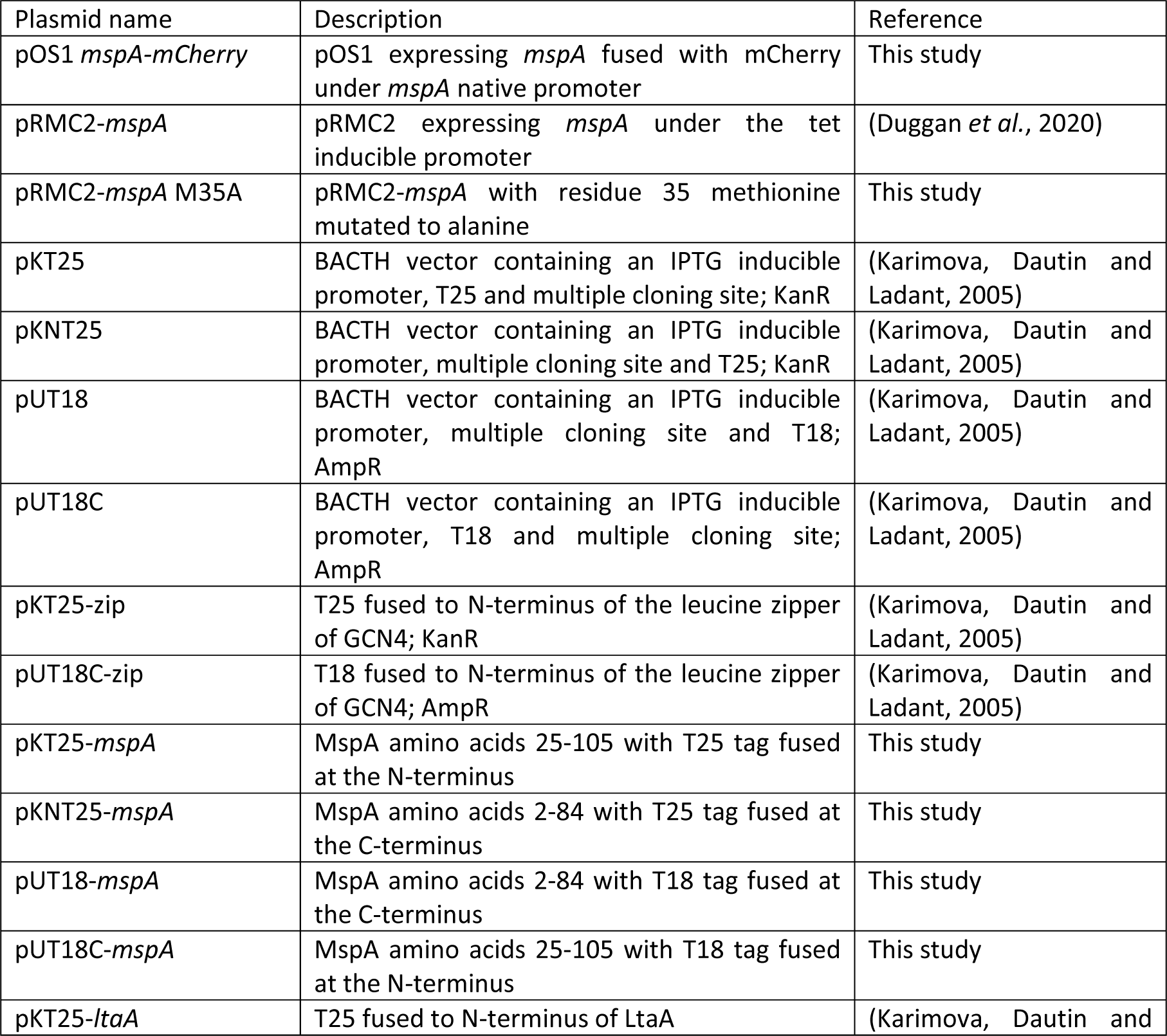
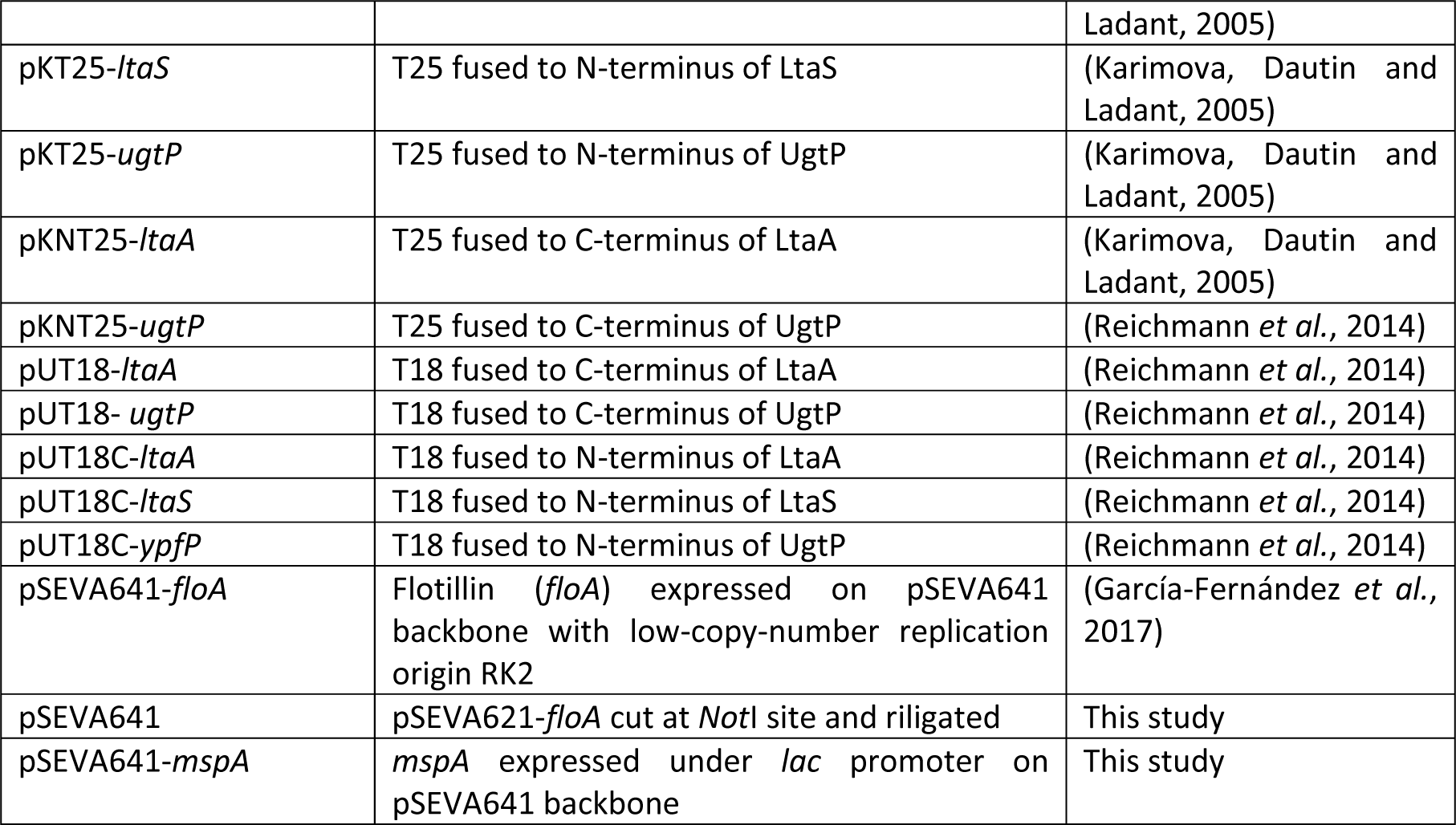
Plasmids used in this study.

**Table S3.**
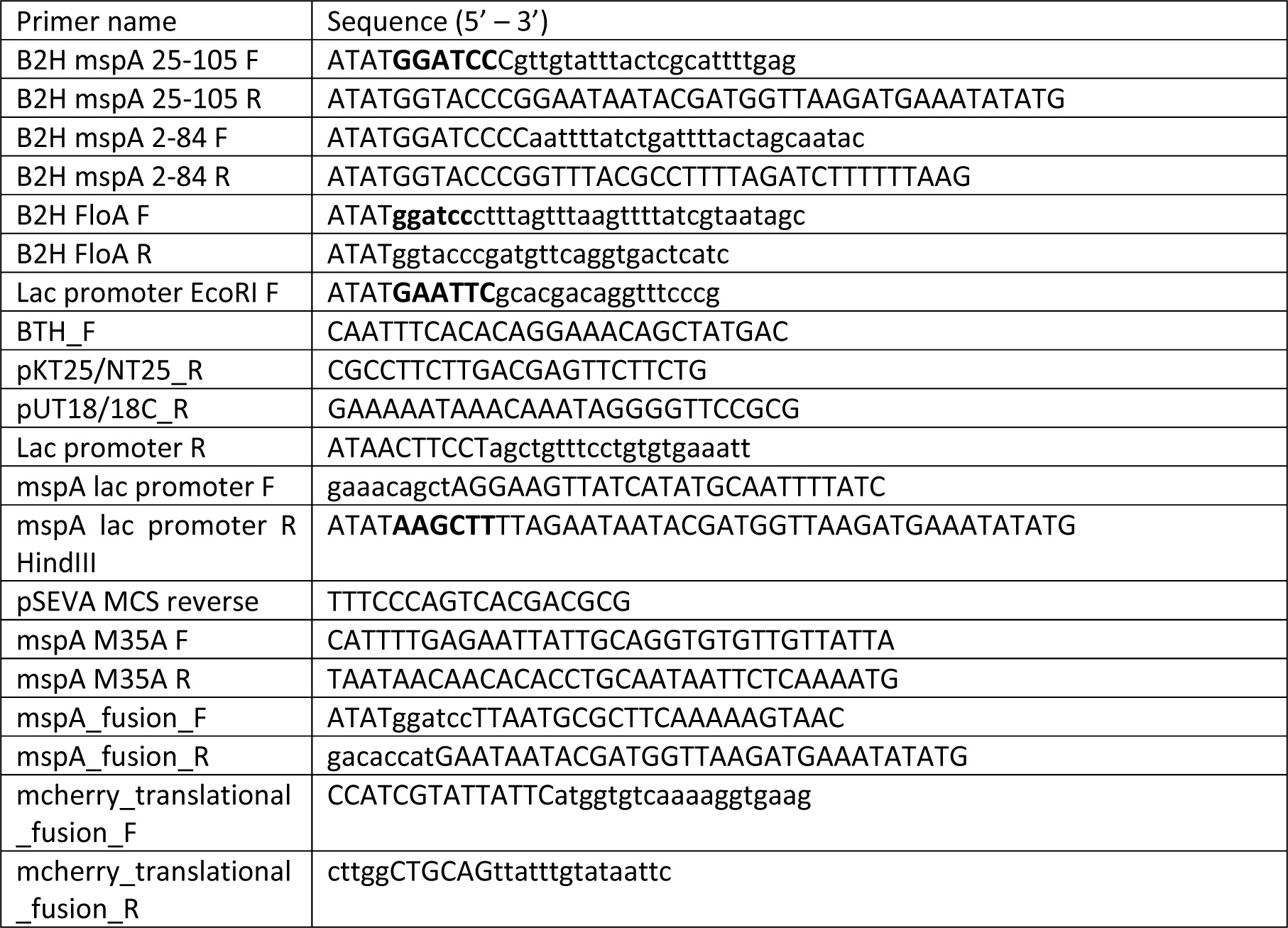
Primers used in this study.

